# LEP-AD: Language Embedding of Proteins and Attention to Drugs predicts drug target interactions

**DOI:** 10.1101/2023.03.14.532563

**Authors:** Anuj Daga, Sumeer Ahmad Khan, David Gomez Cabrero, Robert Hoehndorf, Narsis A. Kiani, Jesper Tegnér

## Abstract

Predicting drug-target interactions is a tremendous challenge for drug development and lead optimization. Recent advances include training algorithms to learn drug-target interactions from data and molecular simulations. Here we utilize Evolutionary Scale Modeling (ESM-2) models to establish a Transformer protein language model for drug-target interaction predictions. Our architecture, LEP-AD, combines pre-trained ESM-2 and Transformer-GCN models predicting binding affinity values. We report new best-in-class state-of-the-art results compared to competing methods such as SimBoost, DeepCPI, Attention-DTA, GraphDTA, and more using multiple datasets, including Davis, KIBA, DTC, Metz, ToxCast, and STITCH. Finally, we find that a pre-trained model with embedding of proteins (the LED-AD) outperforms a model using an explicit alpha-fold 3D representation of proteins (e.g., LEP-AD supervised by Alphafold). The LEP-AD model scales favorably in performance with the size of training data. Code available at https://github.com/adaga06/LEP-AD

## 1 Introduction

Successful drug development requires a solid understanding of the molecular mechanisms underlying the drug’s mechanism of action. One of the most crucial components of this understanding is the identification of Drug-Target Interactions (DTI), which determine if a chemical compound would affect a particular target/protein, called binding affinity. Binding affinity is decisive in determining drug efficacy and safety and identifying specific target proteins.

Despite the importance of identifying DTIs, it remains a challenging task due to the heterogeneous nature of targets, sparsity of data, inter-individual variability, and an incomplete understanding of molecular mechanisms. Investigators have turned to computational methods that integrate molecular modeling, computer simulations, and experimental data to address these challenges. Such approaches strive to gain insights into the molecular details of drug-target interactions, predict binding affinity, and thereby guide the selection of drug candidates before clinical trials.

This paper presents a new approach, “LEP-AD”, where we combine a deep latent embedding of proteins using a language model with a graph-based representation of drugs with attention as computed in a Transformer model.

In this paper, we evaluate the performance of LEP-AD against existing state-of-the-art methods and demonstrate its ability to predict binding affinity between drugs and target proteins accurately. The results of our experiments indicate that LEP-AD provides significant improvements over previous methods in terms of accuracy and speed. Overall, our findings demonstrate the potential of LEP-AD to serve as a valuable tool for drug discovery and development, providing insights into molecular mechanisms of drug-target interactions and guiding the selection of drug candidates for clinical trials.

## 2 Related Work

The prediction of binding affinity, or the binding strength between a drug and its target in the body, is a crucial aspect of drug discovery and development. Accurate binding affinity predictions can help identify the most promising drug candidates, optimize the design of new drugs, and reduce the cost and time of drug development. The high computational cost of methods based on molecular dynamics and quantum mechanics, despite the reliable results, limits their use in high-throughput screening. And those based on molecular docking have faster computation suffering from low accuracy. A growing number of researchers have favored deep learning methods as they provide a better trade-off between computational cost and accuracy Nguyen et al. (2019) Öüzturk et al. (2018).

Traditional machine learning methods, such as decision trees, random forests, and support vector machines, have also been widely used for binding affinity predictions in the past Kinnings et al. (2011) Iskar et al. (2012) Corsello et al. (2017). However, these methods rely on hand-crafted features and have limited ability to capture the complex and diverse relationships between drugs and targets. The end-to-end deep learning based methods Tsubaki et al. (2019b) are based on artificial neural networks, which are designed to automatically learn the complex relationships between drugs and targets from large amounts of data. Deep learning approaches for binding affinity prediction include convolutional neural networks (CNNs) Öztürk et al. (2018), long short-term memory networks (LSTMs) Abbasi et al. (2020) Mukherjee et al., self-attention mechanisms Shin et al. (2019b), and generative adversarial networks (GANs) Zhao et al. (2020).

Despite the advantages of deep learning-based methods, they also have some limitations. One of the major limitations is that these methods typically convert drug compounds into string representations, such as SMILES (Simplified Molecular Input Line Entry System) strings, which do not preserve the topological information of the drug molecules. The function of proteins greatly depends on their tertiary structures and, consequently, the binding affinity – for example, the size, characteristics, and number of protein pockets and cavities can affect the binding significantly. This can lead to the loss of important structural information and result in decreased accuracy in binding affinity predictions. Not having access to many proteins with structural information has also been an obstacle to developing such methods. After introducing the concept of graph neural network (GNN) researchers start using it for predicting DTIs because of its impressive performance, high interpretability and the fact that GNNs are generally less sensitive to the choice of atomic descriptors, unlike traditional feature engineering-based ML models Nguyen et al. (2019). Graph neural networks can capture the topological information of drug molecules and can be seen as a way to capture important structure information with reasonable computational cost in contrast to other architectures such as CNN. In these Structurally aware GNN methods such as GraphDTA Nguyen et al. (2019), the chemical structures of drugs are represented as graphs, where atoms are represented as nodes, interactions between atoms are represented as edges, and proteins are represented as strings. Graph neural networks, such as graph convolutional networks (GCNs), graph attention networks (GATs), and graph isomorphism networks (GINs), are then applied to predict binding affinity. These methods have shown promising results in improving the accuracy of binding affinity predictions by incorporating the topological information of drug molecules.

In this study, we present a novel approach, LEP-AD, which takes into account for the latent structure of proteins for the prediction of drug-target interactions. Our approach has been shown to significantly improve results compared to the state-of-the-art methods. Since previous studies have attempted to utilize pretrained Alphafold and Evolutionary Sequence Model (ESM) approaches Kalakoti et al. (2022), these earlier and smaller models have not been able to match the current state-of-the-art performance in the prediction of drug-target interactions. To this end, we also assessed the performance of LEP-AD model supervised by the Alphafold model, thus taking the tertiary protein structure into account. We compare our results with KronRLS Cichonska et al. (2017), Cichonska et al. (2018), SimBoost He et al. (2017), DeepDTA Öztürk et al. (2018), Mt-Dti Shin et al. (2019a), DeepCPI Tsubaki et al. (2019a), WideDTA Öztürk et al. (2019), GANsDTA Zhao et al. (2020), AttentionDTA Zhao et al. (2019), 1D-CNN Majumdar et al. (2021), DeepGS Lin (2020), and GraphDTA Nguyen et al. (2019).

## 3. Method

The proposed LEP-AD encoder-decoder architecture is designed to predict protein-ligand binding affinity, as shown in Figure 1. The encoding part of the model consists of two modules. The first module extracts topological information from drug molecules, while the second module extracts sequential information from target proteins. To prepare protein sequences for machine learning algorithms, we convert them into a numerical representation by transforming the protein sequence into a numerical form. We explored the intuition that the tertiary protein structures would be advantageous for our prediction problem. To this end, we utilized the ESM-2 Lin et al. (2022) pre-trained model with 3B parameters. We also have implemented and used methods such as Alphafold-2, Openfold, and Fastfold on a smaller dataset. We refer to this model as LEP-AD supervised by Alphafold.

**Figure 1:**
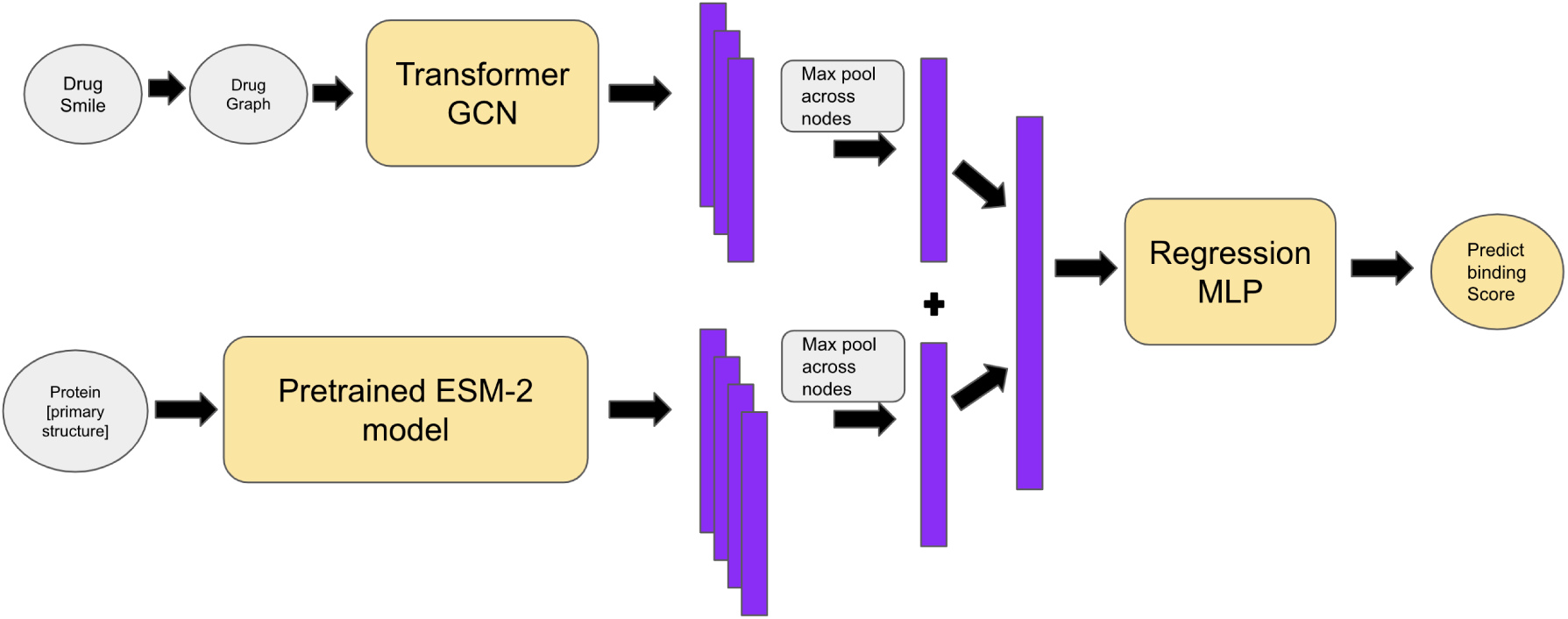
LEP-AD Model Architecture

**Figure 2:**
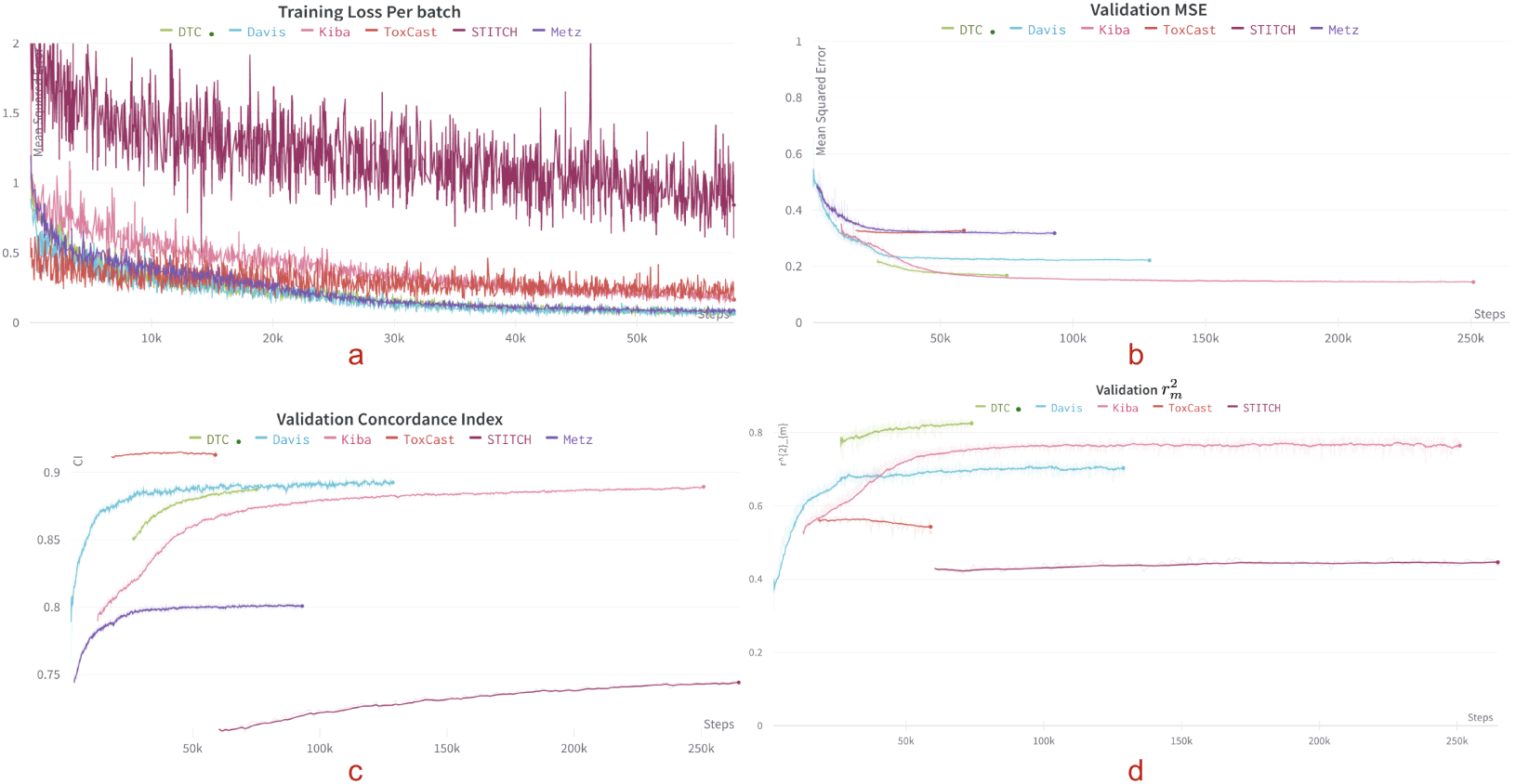
a.) Training loss per batch. b.)Validation loss(MSE) to ensure that the model is not over-fitting. c.) Concordance Index (CI) plot as an evaluation metric on validation data. d.) 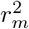plot onvalidation data.

**Figure 3:**
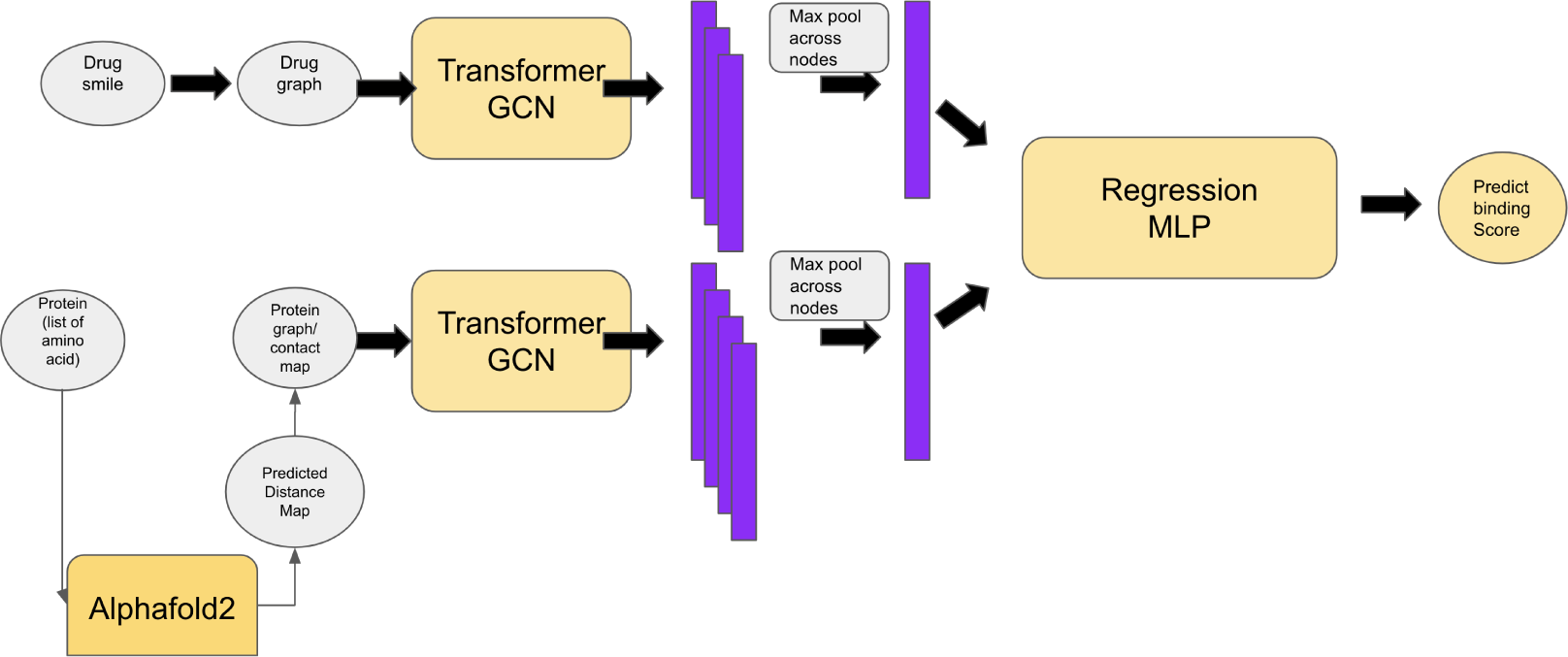
LEP-AD-Variant with supervised Alphafold2 model

**Figure 4:**
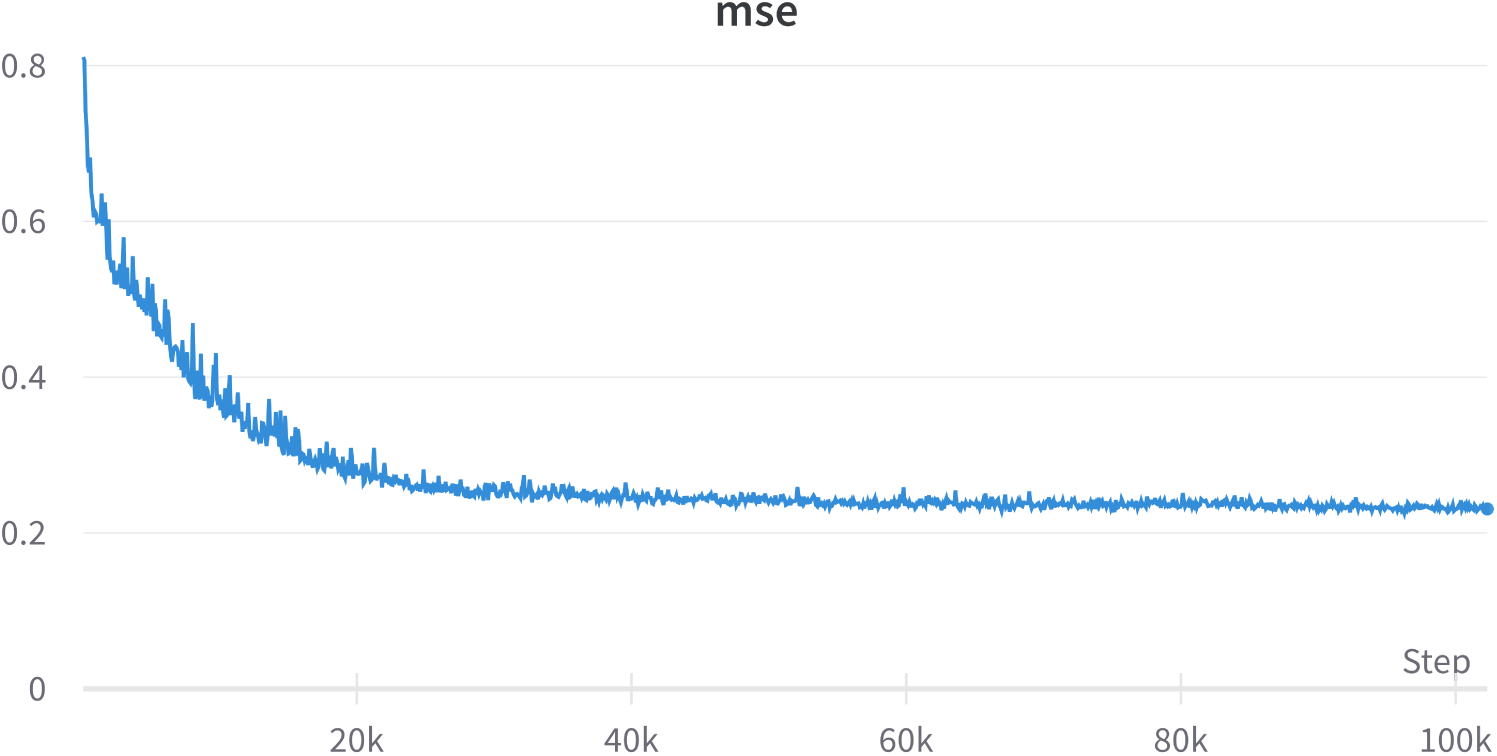
Validation Loss(MSE) plot of LEP-AD-Variant with supervised Alphafold2 model

**Figure 5:**
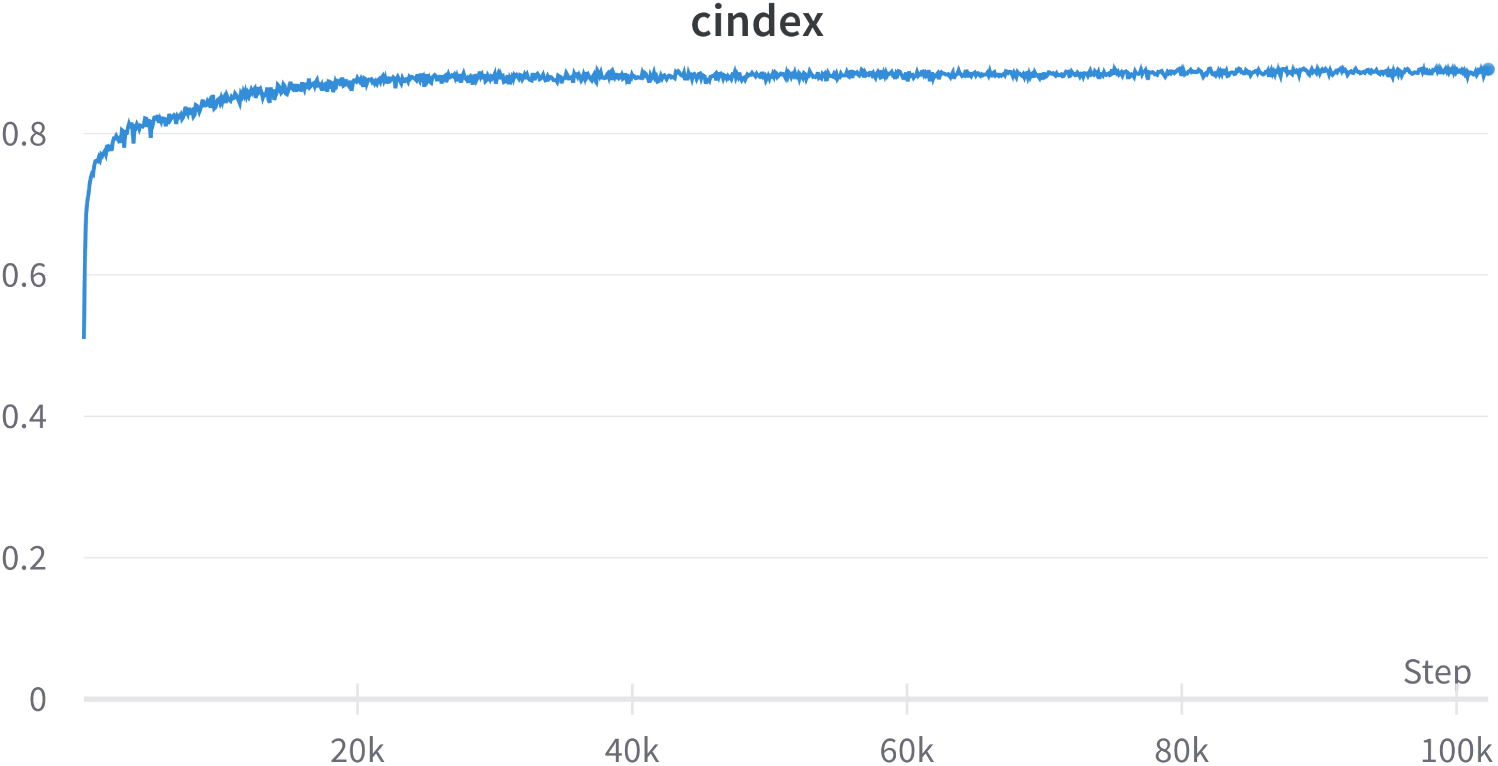
CI plot of LEP-AD-Variant with supervised Alphafold2 model

The ESM-2 model generates latent representations of each amino acid in the protein sequence, thus yielding a compact, non-3D representation of the entire protein sequence through a global max pooling operation. To capture the topological information of drug compounds, each drug/small compound was represented using its Simplified Molecular Input Line Entry System (SMILES) notation. This is converted into a graph representation using open-source chemical informatics software RD-Kit. We apply GCN Kipf & Welling (2016) and Transformer Vaswani et al. (2017) calculations to the attention matrix in the Graph Transformer Shi et al. (2020). Propagating features from one layer to the next GCN layers follows:

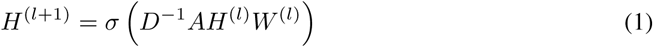

The Transformer equation for calculating attention across the value vector is 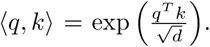 Applying these GCN propagating layers to Transformers, we have Shi et al. (2020) the attention matrix from their Graph Transformer is:

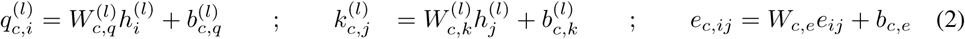

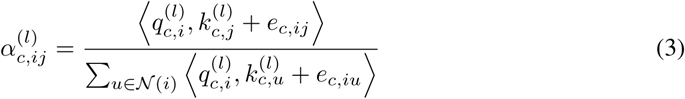

Their model Shi et al. (2020) unifies both label propagation and feature propagation within a shared message-passing framework. This framework Shi et al. (2020) incorporates attention mechanism into the graph neural network layers using scalar dot product attention across the value vector

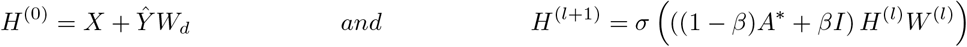

This second module captures the topological information about the drug by applying self-attention mechanisms over the nodes of the graph representation of the SMILES notation. The output from both modules is concatenated and fed into a fully connected layer to predict the binding affinity between the drug and target.

It is noteworthy that, Initially, the Graph Transformer model was selected as the state-of-the-art model for both drugs and targets. However, it became apparent that targets are significantly longer sequences compared to drugs, resulting in high computational complexity and sensitivity during training. Consequently, we decided to utilize the pretrained ESM-2 model, which has been trained on a vast dataset, to significantly reduce computational time while maintaining high accuracy.

The decoder part of the model is a regression model, with three layers that take the protein and drug representations as inputs and predict the binding affinity.

In summary, the LEP-AD architecture captures crucial information for drug-target interaction prediction by extracting topological information from drug molecules and sequential information from target proteins. The combination of a graph representation of the SMILES notation and Transformer-based GCN layers enhances the model’s performance and produces more accurate results.

## 4. Experiments

### 4.1 Dataset

We used Davis Davis MI (2011), KIBA Tang et al. (2014), Drug Target Commons (DTC) Tang (2018), Metz Metz (2011), ToxCast Tox and STITCH Kuhn (2007) datasets for our experiments as summarized in Table 1.

**Table 1:** Dataset statistics

| Dataset | # of drugs<br>(compounds) | # of targets<br>proteins | Total number of<br>drug-target pairs used |
| --- | --- | --- | --- |
| Davis ($k_d$) | 68 | 442 | 30056 |
| KIBA | 2111 | 229 | 118254 |
| DTC ($pk_i$) | 5983 | 118 | 67894 |
| Metz ($pk_i$) | 1471 | 170 | 35307 |
| ToxCast ($AC_{50}$) | 7657 | 328 | 342869 |
| STITCH | 724471 | 15258 | 1244420 |

### 4.2 Baselines

We used GraphDTA Nguyen et al. (2019) as a baseline model. Additional baseline models include KronRLS Cichonska et al. (2017), Cichonska et al. (2018), SimBoost He et al. (2017), DeepDTA Öztürk et al. (2018), Mt-Dti Shin et al. (2019a), DeepCPI Tsubaki et al. (2019a), WideDTA Öztürk et al. (2019), GANsDTA Zhao et al. (2020), AttentionDTA Zhao et al. (2019), 1D-CNN Majumdar et al. (2021), DeepGS Lin (2020).

### 4.3 Evaluation Metric

MSE is a commonly used metric to measure the difference between the predicted value and the real value. For n samples, the MSE is calculated as the average of the sum of the square of the difference between the predicted value p (i = 1, 2,…, n) and the real value y. A smaller MSE means that the predicted values of the sample are closer to the real values.

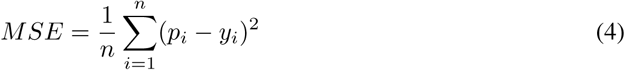

We use the *r*^2^ metric 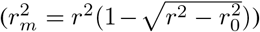 Roy et al. (2013) to evaluate prediction performance of Quantitative Structure-Activity Relationship (QSAR) models. The predictions of a QSAR model are deemed acceptable if its *r*^2^ value is equal to or greater than 0.5. Here *r*^2^ and 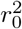 and the squared correlation coefficient values between the ground truth and the predicted values with and without intercept, respectively.

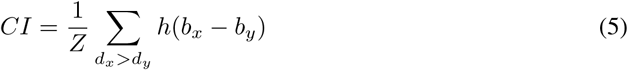

CI measures the distinction between the predicted value and the real value in the analysis, calculated as

### 4.4 Hyperparameters

To ensure fairness in our comparison, we employed the same set of training and testing examples from the Nguyen et al. (2019) database, as well as the same performance metrics - Mean Square Error (MSE, where lower values are preferred) and Concordance Index and 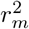 (where higher values are preferred). For the baseline methods, we present the performance metrics as reported in the Nguyen et al. (2019) database. The hyper-parameters utilized in our experiments are outlined in Table 2, and were selected a priori based on our previous modeling experience without any further tuning.

**Table 2:** Hyper-parameters used in our experiments

| Hyperparameter | Settings |
| --- | --- |
| Learning rate | 0.0005 |
| Batch Size | 512 |
| Optimizer | Adam |
| Loss | MSE |
| Dropout | 0.2 |
| Transformer GCN Layers[Drug] | [8 heads(78-78),4 heads(624-156),2 heads(624-312)] |
| Linear Layers[Drug] | [624,1024,128] |
| ESM-2 dimensions [3B parameters,36 layers] | 2560 |
| Linear Layers | [2688,1024,512,1] |

## 5 Results

We evaluate the performance of all models across all datasets reported in Tables 3,4 and 5 using the concordance index (CI), 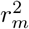 index, and mean squared error (MSE). Additionally, we evaluated the impact of replacing the pretrained ESM model with Alphafold2 on the performance of our predictive model. While Alphafold2 (LEP-AD supervised by Alphafold) demonstrated improved accuracy in predicting protein structures compared to the current state of the art, the improvement was competitive compared to the previous model. More details about this model are in Appendix. Interestingly, the pretrained ESM model yielded an additional improvement (Table 3). Furthermore, the process of generating multiple sequence alignments (MSA) using Alphafold2 resulted in slower processing times. The results of our evaluation are presented in Table 3, which were obtained using the Davis dataset. Despite the potential for better performance with Alphafold2, we ultimately chose to retrain the pretrained ESM-2 model for our predictive model due to its higher speed, better scalability, and better overall performance.

**Table 3:** Performance comparison of LEP-AD against all other baseline models using Davis (pKd) and KIBA data

| Model | Davis (pK<sub>d</sub>) |  |  | KIBA |  |  |
| --- | --- | --- | --- | --- | --- | --- |
|  | MSE | CI | r<sub>m</sub><sup>2</sup> | MSE | CI | r<sub>m</sub><sup>2</sup> |
| KronRLS | 0.379 | 0.869 | 0.407 | 0.411 | 0.782 | 0.342 |
| SimBoost | 0.282 | 0.873 | 0.644 | 0.222 | 0.836 | 0.629 |
| DeepDTA | 0.261 | 0.878 | 0.630 | 0.194 | 0.863 | 0.673 |
| MT-DTI (Wo-FT) | 0.268 | 0.875 | 0.633 | 0.220 | 0.844 | 0.584 |
| MT-DTI | 0.245 | 0.887 | 0.665 | 0.193 | 0.882 | 0.738 |
| DeepCPI | 0.293 | 0.867 | 0.607 | 0.211 | 0.852 | 0.657 |
| WideDTA | 0.262 | 0.886 | 0.633 | 0.179 | 0.875 | 0.675 |
| GANsDTA | 0.276 | 0.881 | 0.653 | 0.224 | 0.866 | 0.775 |
| Attention-DTA | 0.245 | 0.887 | 0.657 | 0.162 | 0.882 | 0.735 |
| 1D-CNN | — | — | — | 0.700 | — | — |
| DeepGS | 0.252 | 0.880 | 0.686 | 0.193 | 0.860 | 0.684 |
| GraphDTA (GCN) | 0.254 | 0.880 | — | 0.139 | 0.889 | — |
| GraphDTA (GAT_GCN) | 0.245 | 0.881 | — | 0.139 | 0.891 | — |
| GraphDTA (GAT) | 0.232 | 0.892 | 0.662 | 0.179 | 0.866 | 0.671 |
| GraphDTA (GIN) | 0.229 | 0.893 | 0.649 | 0.147 | 0.882 | 0.684 |
| LEP-AD(supervised by Alphafold) | **0.228** | **0.894** | **0.692** | **0.133** | **0.893** | **0.752** |
| LEP-AD* | **0.222** | **0.896** | **0.723** | **0.135** | **0.895** | **0.780** |

**Table 4:** Performance comparison of LEP-AD against all other baseline models on data from DTC (*pk*_*i*_) and Metz (*pk*_*i*_)

| Model | DTC ($pk_i$) | | | Metz ($pk_i$) | | |
| --- | --- | --- | --- | --- | --- | --- |
|  | MSE | CI | r<sub>m</sub><sup>2</sup> | MSE | CI | r<sub>m</sub><sup>2</sup> |
| GraphDTA (GCN) | 0.317 | 0.878 | 0.812 | 0.317 | 0.801 | 0.620 |
| GraphDTA (GAT_GCN) | 0.200 | 0.857 | 0.790 | 0.333 | 0.795 | 0.602 |
| GraphDTA (GAT) | 0.195 | 0.859 | 0.788 | 0.393 | 0.775 | 0.549 |
| GraphDTA (GIN) | 0.176 | 0.876 | 0.798 | 0.317 | 0.800 | 0.645 |
| LEP-AD* | **0.171** | **0.881** | **0.830** | **0.292** | **0.810** | **0.682** |

**Table 5:** Performance comparison LEP-AD of against all other baseline models on data from ToxCast (*pk*_*i*_) and STITCH (*pk*_*i*_)

| Model | ToxCast ($AC_{50}$) | | | STITCH | | |
| --- | --- | --- | --- | --- | --- | --- |
|  | MSE | CI | r<sub>m</sub><sup>2</sup> | MSE | CI | r<sub>m</sub><sup>2</sup> |
| GraphDTA (GCN) | **0.316** | 0.917 | 0.561 | 1.022 | — | 0.424 |
| GraphDTA (GAT_GCN) | 0.317 | 0.917 | 0.566 | — | — | — |
| GraphDTA (GAT) | 0.346 | 0.911 | 0.535 | — | — | — |
| GraphDTA (GIN) | 0.324 | 0.915 | 0.555 | — | — | — |
| LEP-AD* | **0.316** | **0.918** | **0.572** | **0.986** | **0.745** | **0.461** |

It is noteworthy that, even in larger datasets, there was a considerable enhancement in the squared correlation coefficient (also referred to as the coefficient of determination, or 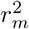) metric, while the mean squared error (MSE) and concordance index showed less improvement, while 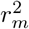shows a clearincreasing trend in all the datasets mentioned above. Our method showed significant improvement in the R2 metric, with an increase of around 2% for the ToxCast and DTC, around 4% for the STITCH dataset and Metz, and around 5% for the Davis datasets compared to the second-best method. These results suggest that our method could be a strong candidate for the state-of-the-art model predicting binding affinity.

This disparity in improvement between the *r*^2^ and other metrics could be due to a stronger linear relationship between the ground truth and predicted values contributing to the enhancement in *r*^2^, while the MSE and concordance index is more sensitive to factors such as outliers or factors not captured by the linear relationship between the ground-truth and predicted values.

## 6. Conclusion

Here we demonstrate the potential of using a sizeable pre-trained protein language model and modeling as a comprehensive approach to uncover molecular mechanisms underlying drug-target interactions. Interestingly, using a pretrained model appeared to be advantageous compared to explicitly using a 3D protein representation. This observation is also advantageous since using a pretrained embedding approach scales, as of now, better compared to computationally intensive 3D calculation. It remains to be investigated whether using a large library of precomputed 3D structures could be competitive with a pretrained embedding strategy. By analyzing larger datasets and validating our results through additional experiments, we aim to improve our method further and gain a deeper understanding of these mechanisms. In particular, our study reveals that there is a need to dissect the relationship between the size of the data and the performance when different metrics are used. Specifically, an analysis where methods fail using additional metrics could provide clues for further enhancement of our DTI method.

Our results have already shown new state-of-the-art performance using several key metrics and data sets. Accurately predicting drug-target interactions can revolutionize drug development by leading to the selection of more promising drug candidates and the optimization of their properties before clinical trials. This, in turn, can significantly improve the success rate of drug development and result in the discovery of new, more effective, and safer drugs.

In summary, we believe that our future work will continue to shape the future of drug development and improve human health by discovering new and effective treatments for a wide range of diseases. Our results demonstrate our method’s potential to impact this field significantly and ultimately benefit human health.

## A Supplementary

### Methods

The novel aspect of this model lies in the encoding of proteins. To encode the proteins, we employed the Alphafold2 method to generate the predicted distance matrix. From the 442 proteins in the Davis dataset, some were obtained directly from the Alphafold database (https://alphafold.ebi.ac.uk) while others were generated using the method proposed by Ahdritz et al. (2022). We used this implementation with its default parameters. After generating the predicted protein structure, we normalized it and applied a cutoff to convert it into a binary matrix, resulting in a contact map, which is an unweighted and undirected graph adjacency matrix. In a manner similar to the drug graph, we fed the protein contact map into the Transformer-GCN network to obtain the latent representation of each node in the protein contact map.

## Notes

### Competing Interest Statement

The authors have declared no competing interest.

## References

US, EPA summary files from invitrodb v2, accessed on 2018-03-22. URL https://www.epa.gov/chemical-research/toxicity-forecaster-toxcasttm-data.

Karim Abbasi, Parvin Razzaghi, Antti Poso, Massoud Amanlou, Jahan B Ghasemi, and Ali Masoudi-Nejad. DeepCDA: deep cross-domain compoundÄiprotein affinity prediction through LSTM and convolutional neural networks. Bioinformatics, 36(17):4633–4642, November 2020. ISSN 1367-4803, 1460-2059. doi: 10.1093/bioinformatics/btaa544. URL https://academic.oup.com/bioinformatics/article/36/17/4633/5848004.

Gustaf Ahdritz, Nazim Bouatta, Sachin Kadyan, Qinghui Xia, William Gerecke, Timothy J O’Donnell, Daniel Berenberg, Ian Fisk, Niccolò Zanichelli, Bo Zhang, Arkadiusz Nowaczynski, Bei Wang, Marta M Stepniewska-Dziubinska, Shang Zhang, Adegoke Ojewole, Murat Efe Guney, Stella Biderman, Andrew M Watkins, Stephen Ra, Pablo Ribalta Lorenzo, Lucas Nivon, Brian Weitzner, Yih-En Andrew Ban, Peter K Sorger, Emad Mostaque, Zhao Zhang, Richard Bonneau, and Mohammed AlQuraishi. Openfold: Retraining alphafold2 yields new insights into its learning mechanisms and capacity for generalization. bioRxiv, 2022. doi: 10.1101/2022.11.20.517210. URL https://www.biorxiv.org/content/10.1101/2022.11.20.517210.

Anna Cichonska, Balaguru Ravikumar, Elina Parri, Sanna Timonen, Tapio Pahikkala, Antti Airola, Krister Wennerberg, Juho Rousu, and Tero Aittokallio. Computational-experimental approach to drug-target interaction mapping: A case study on kinase inhibitors. PLOS Computational Biology, 13(8):e1005678, August 2017. ISSN 1553-7358. doi: 10.1371/journal.pcbi.1005678. URL https://dx.plos.org/10.1371/journal.pcbi.1005678.

Anna Cichonska, Tapio Pahikkala, Sandor Szedmak, Heli Julkunen, Antti Airola, Markus Heinonen, Tero Aittokallio, and Juho Rousu. Learning with multiple pairwise kernels for drug bioactivity prediction. Bioinformatics, 34(13):i509–i518, July 2018. ISSN 1367-4803, 1460-2059. doi: 10.1093/bioinformatics/bty277. URL https://academic.oup.com/bioinformatics/article/34/13/i509/5045738.

Steven M Corsello, Joshua A Bittker, Zihan Liu, Joshua Gould, Patrick McCarren, Jodi E Hirschman, Stephen E Johnston, Anita Vrcic, Bang Wong, Mariya Khan, Jacob Asiedu, Rajiv Narayan, Christopher C Mader, Aravind Subramanian, and Todd R Golub. The Drug Re-purposing Hub: a next-generation drug library and information resource. Nature Medicine, 23(4):405–408, April 2017. ISSN 1078-8956, 1546-170X. doi: 10.1038/nm.4306. URL http://www.nature.com/articles/nm.4306.

Herrgard S et al. Davis MI, Hunt JP. Comprehensive analysis of kinase inhibitor selectivity. Nature biotechnology, 29(11), 2011. doi: 10.1038/nbt.1990.

Tong He, Marten Heidemeyer, Fuqiang Ban, Artem Cherkasov, and Martin Ester. SimBoost: a read-across approach for predicting drugÄ ‘itarget binding affinities using gradient boosting machines. Journal of Cheminformatics, 9(1):24, December 2017. ISSN 1758-2946. doi: 10.1186/s13321-017-0209-z. URL https://jcheminf.biomedcentral.com/articles/10.1186/s13321-017-0209-z.

Murat Iskar, Georg Zeller, Xing-Ming Zhao, Vera van Noort, and Peer Bork. Drug discovery in the age of systems biology: the rise of computational approaches for data integration. Current Opinion in Biotechnology, 23(4):609–616, August 2012. ISSN 09581669. doi: 10.1016/j.copbio.2011.11.010. URL https://linkinghub.elsevier.com/retrieve/pii/ S095816691100721X.

Yogesh Kalakoti, Shashank Yadav, and Durai Sundar. TransDTI: Transformer-Based Language Models for Estimating DTIs and Building a Drug Recommendation Workflow. ACS Omega, 7 (3):2706–2717, January 2022. ISSN 2470-1343, 2470-1343. doi: 10.1021/acsomega.1c05203. URL https://pubs.acs.org/doi/10.1021/acsomega.1c05203.

Sarah L. Kinnings, Nina Liu, Peter J. Tonge, Richard M. Jackson, Lei Xie, and Philip E. Bourne. A Machine Learning-Based Method To Improve Docking Scoring Functions and Its Application to Drug Repurposing. Journal of Chemical Information and Modeling, 51(2):408–419, February 2011. ISSN 1549-9596, 1549-960X. doi: 10.1021/ci100369f. URL https://pubs.acs. org/doi/10.1021/ci100369f.

Thomas N. Kipf and Max Welling. Semi-supervised classification with graph convolutional networks, 2016. URL https://arxiv.org/abs/1609.02907.

Mering C. Campillos M. Jensen L. Bork P. Kuhn, M. Stitch: interaction networks of chemicals and proteins. Nucleic Acids Research, 36, 2007.

Xuan Lin. DeepGS: Deep Representation Learning of Graphs and Sequences for Drug-Target Binding Affinity Prediction, April 2020. URL http://arxiv.org/abs/2003.13902. arXiv:2003.13902 [cs, q-bio].

Zeming Lin, Halil Akin, Roshan Rao, Brian Hie, Zhongkai Zhu, Wenting Lu, Nikita Smetanin, Robert Verkuil, Ori Kabeli, Yaniv Shmueli, Allan dos Santos Costa, Maryam Fazel-Zarandi, Tom Sercu, Salvatore Candido, and Alexander Rives. Evolutionary-scale prediction of atomic level protein structure with a language model. bioRxiv, 2022. doi: 10.1101/2022.07.20.500902. URL https://www.biorxiv.org/content/early/2022/12/21/2022.07.20.500902.

Shatadru Majumdar, Soumik Kumar Nandi, Shuvam Ghosal, Bavrabi Ghosh, Writam Mallik, Nilanjana Dutta Roy, Arindam Biswas, Subhankar Mukherjee, Souvik Pal, and Nabarun Bhattacharyya. Deep Learning-Based Potential Ligand Prediction Framework for COVID-19 with DrugÄ ‘iTarget Interaction Model. Cognitive Computation, February 2021. ISSN 1866-9956, 1866-9964. doi: 10.1007/s12559-021-09840-x. URL http://link.springer.com/10.1007/s12559-021-09840-x.

Johnson E. Soni N. Merta P. Kifle L. Hajduk -P. Metz, J. Navigating the kinome. Nature Chemical-Biology, 7, 2011.

Shrimon Mukherjee, Madhusudan Ghosh, and Partha Basuchowdhuri. DeepGLSTM: Deep Graph Convolutional Network and LSTM based approach for predicting drug-target binding affinity, pp. 729–737. doi: 10.1137/1.9781611977172.82. URL https://epubs.siam.org/doi/abs/10.1137/1.9781611977172.82.

Thin Nguyen, Hang Le, and Svetha Venkatesh. Graphdta: prediction of drug–target binding affinity using graph convolutional networks. bioRxiv, 2019. doi: 10.1101/684662.

Kunal Roy, Pratim Chakraborty, Indrani Mitra, Probir Kumar Ojha, Supratik Kar, and Rudra Narayan Das. Some case studies on application of Ä úrm2Äu‘ metrics for judging quality of quantitative structureÄ‘iactivity relationship predictions: Emphasis on scaling of response data. Journal of Computational Chemistry, 34(12):1071–1082, 2013. doi: https://doi.org/10.1002/jcc.23231. URL https://onlinelibrary.wiley.com/doi/abs/10.1002/jcc.23231.

Yunsheng Shi, Zhengjie Huang, Shikun Feng, Hui Zhong, Wenjin Wang, and Yu Sun. Masked label prediction: Unified message passing model for semi-supervised classification, 2020. URL https://arxiv.org/abs/2009.03509.

Bonggun Shin, Sungsoo Park, Keunsoo Kang, and Joyce C. Ho. Self-attention based molecule representation for predicting drug-target interaction. In Finale Doshi-Velez, Jim Fackler, Ken Jung, David Kale, Rajesh Ranganath, Byron Wallace, and Jenna Wiens (eds.), Proceedings of the 4th Machine Learning for Healthcare Conference, volume 106 of Proceedings of Machine Learning Research, pp. 230–248. PMLR, 09–10 Aug 2019a. URL https://proceedings.mlr.press/v106/shin19a.html.

Bonggun Shin, Sungsoo Park, Keunsoo Kang, and Joyce C. Ho. Self-attention based molecule representation for predicting drug-target interaction, 2019b. URL https://arxiv.org/abs/1908.06760.

Jing Tang, Agnieszka Szwajda, Sushil Shakyawar, Tao Xu, Petteri Hintsanen, Krister Wennerberg, and Tero Aittokallio. Making sense of large-scale kinase inhibitor bioactivity data sets: A comparative and integrative analysis. Journal of Chemical Information and Modeling, 54, 2014. doi: 10.1021/ci400709d.

Ravikumar B. Alam Z. Rebane A. V ah a-Koskela M. Peddinti G. Adrichem A. Wakkinen J. Jaiswal A. Karjalainen Tang, Jing. Others drug target com-mons: a community effort to build a consensus knowl-edge base for drug-target interactions. Cell Chemical Biology, 25, 2018. doi: 10.1021/ci400709d.

Masashi Tsubaki, Kentaro Tomii, and Jun Sese. CompoundÄ ‘iprotein interaction prediction with end-to-end learning of neural networks for graphs and sequences. Bioinformatics, 35(2):309– 318, January 2019a. ISSN 1367-4803, 1367-4811. doi: 10.1093/bioinformatics/bty535. URL https://academic.oup.com/bioinformatics/article/35/2/309/5050020.

Masashi Tsubaki, Kentaro Tomii, and Jun Sese. CompoundÄ ‘iprotein interaction prediction with end-to-end learning of neural networks for graphs and sequences. Bioinformatics, 35(2):309– 318, January 2019b. ISSN 1367-4803, 1367-4811. doi: 10.1093/bioinformatics/bty535. URL https://academic.oup.com/bioinformatics/article/35/2/309/5050020.

Ashish Vaswani, Noam Shazeer, Niki Parmar, Jakob Uszkoreit, Llion Jones, Aidan N. Gomez, Lukasz Kaiser, and Illia Polosukhin. Attention is all you need, 2017. URL https://arxiv. org/abs/1706.03762.

Lingling Zhao, Junjie Wang, Long Pang, Yang Liu, and Jun Zhang. Gansdta: Predicting drug-target binding affinity using gans. Frontiers in Genetics, 10, 2020. ISSN 1664-8021. doi: 10.3389/fgene.2019.01243. URL https://www.frontiersin.org/articles/10.3389/fgene.2019.01243.

Qichang Zhao, Fen Xiao, Mengyun Yang, Yaohang Li, and Jianxin Wang. AttentionDTA: predic-tion of drugÄ‘itarget binding affinity using attention model. In 2019 IEEE International Conference on Bioinformatics and Biomedicine (BIBM), pp. 64–69, San Diego, CA, USA, November 2019. IEEE. ISBN 9781728118673. doi: 10.1109/BIBM47256.2019.8983125. URL https://ieeexplore.ieee.org/document/8983125/.

Hakime Ö ztürk, Arzucan Özgür, and Elif Ozkirimli. DeepDTA: deep drug–target binding affinity prediction. Bioinformatics, 34(17):i821–i829, 09 2018. ISSN 1367-4803. doi: 10.1093/bioinformatics/bty593. URL https://doi.org/10.1093/bioinformatics/bty593.

Hakime Öztürk, Elif Ozkirimli, and Arzucan Ö zgür. Widedta: prediction of drug-target binding affinity, 2019. URL https://arxiv.org/abs/1902.04166.

